# Homology detection using a protein secondary structure-based large language model

**DOI:** 10.1101/2023.12.19.572443

**Authors:** Roman Kogay, Weicheng Ma, Jad Bousselham, Zechen Yang, Daniel Rockmore, Olga Zhaxybayeva, Soroush Vosoughi

**Affiliations:** Department of Biological Sciences, Dartmouth College, Hanover NH, USA; Department of Computer Science, Dartmouth College, Hanover NH, USA; Department of Mathematics, Dartmouth College, Hanover NH, USA; The Santa Fe Institute, Santa Fe, NM, USA

## Abstract

Detection of homology among proteins is fundamental to understanding protein function. Unfortunately, traditional homology searches using amino acid sequence similarity are limited when numerous amino acid substitutions have accumulated either due to billions of years of evolution or through processes of accelerated change. Recent applications of deep-learning approaches demonstrate that “protein language” models of amino acid sequences can improve the accuracy of the traditional homology searches. Ultimately, the ability to work seamlessly with tertiary structures of proteins will solve the homology detection challenge and provide accompanying insights directly related to function, but to date the use of 3D structures suffers both from data availability and computational bottlenecks. Herein, we present the Protein Secondary Structure Language (ProSSL) model, an efficient encoding of protein secondary structure information in a Transformer-based deep-learning architecture. We conjecture that the secondary protein structure, which is better conserved than primary sequences and much more easily predictable and available than tertiary protein structure, could aid in the task of homology detection. ProSSL has the computational advantages of primary sequence-based homology detection, while also providing important structural information for similarity scoring. Using two case studies of large, diverse viral protein families, we show that the ProSSL model successfully captures patterns of secondary structure arrangements and is effective in detecting homologs either as a pre-trained or fine-tuned model. In both tasks, we accurately detect members of these protein families, including those missed in traditional amino acid similarity searches. We also illustrate how functional insights from the individual ProSSL models could be obtained from the use of the Shapley Additive exPlanations (SHAP) values.

**Author Summary:** When DNA is obtained from an organism or an environment, scientists are tasked with determining the functions of the proteins encoded in the genetic material. Such “functional annotation” relies on assigning functions based on the similarity of the proteins to counterparts in databases comprising annotated sequence data. Especially challenging is an ability to recognize similarity in proteins that accumulated a lot of amino acid changes. It is well-known that spatial structure of proteins that share ancestry and perform similar functions evolves much slower than the sequences of the proteins’ amino acids. Thus, comparison of 3D structures could address this challenge, but the data is still limited to certain classes of proteins, and the requisite computations are expensive. Herein we present a deep-learning model derived from protein secondary structure representation, a symbolic encoding of the way neighboring amino acid residues of a protein interact with each other. Unlike 3D structure, the secondary structure is quickly and accurately predictable from amino acid sequences of proteins. Using two viral proteins as case studies, we demonstrate that our model works well for detection of protein similarity, including identification of very distantly related proteins.

## Introduction

Advances in sequencing technologies have substantially accelerated the generation of genomic data, currently acquired at the rate of several exabytes per year [1]. This has dramatically extended the known universe of genes and the proteins encoded by them. A continuing challenge is to understand the functionality of this ever-expanding set of DNA and amino acid sequences. Given that all living organisms share a common ancestor [2], functionality of an unknown sequence can be linked to a known (i.e., functionally annotated) co-descendant or *homolog* [3], identified through *homology search*. The core of homology search is the computation of similarity scores between DNA or amino acid sequences using empirically-derived substitution matrices and fast heuristic algorithms such as BLAST [4] or profile hidden Markov models [5]. Although efficient, such methods are limited in applicability by an inability to establish homology in cases when a large number of nucleotide or amino acid changes have been acquired since the initial divergence of co-descendants, a phenomenon effecting low similarity scores [6]. Distant homologs are common across the Tree of Life and thus remain unrecognized using these techniques. This lacuna is even more pronounced for viruses, whose genetic material is generally subject to higher mutation rates [7]. Downstream effects of such missing information include the incomplete annotation of genomes and metagenomes, in which about 30% of proteins have unknown function [8]. Thus, unrecognized homology adds a significant obstacle to advancing our understanding of biological systems and their evolution.

While primary amino acid sequences may not be usable for homology identification in distantly related proteins, the preservation of 3D (tertiary) structure in proteins, even among homologs with a very low amino acid similarity [6], suggests that tertiary structure comparisons could provide an excellent approach to the problem. However, as of November 2023, the protein tertiary structure databases remain relatively small (∼200,000 experimentally determined structures [9] and ∼200 million AlphaFold models [10]) in comparison to ∼1.2 billion amino acid sequences in the Protein database of GenBank [11]. This size discrepancy is largely due to the high cost and technical challenges associated with both the experiments and computational predictions of the protein tertiary structure. The protein structure databases are also biased towards proteins whose 3D structures are amenable to determination. As a result, homologs will be missed regardless of the efficiency of the tools currently available to compute similarity scores based on tertiary structure features (e.g., DALI [12, 13], TM-ALIGN [14], and even the much more efficient FoldSeek [15]). Indeed, the recent large-scale clustering of protein structures has produced more than 13 million clusters of singletons, which also, on average, have low confidence scores of their AlphaFold structure predictions [16].

Protein secondary structures stand “between” primary and tertiary structures but to date, their potential for homology identification remains largely uncharted. Unlike 3D structure, secondary structure of a protein can be quickly and accurately predicted from the amino acid sequences [17]. The secondary protein structures also remain generally conserved across time, increasing the likelihood that their comparison is attuned to the identification of distant homologs [18]. Finally, the secondary structure is easily represented as sequences drawn from a three-symbol alphabet. With this in mind, we present a Protein Secondary Structure Language (ProSSL) model, which uses the Transformer-based neural network architecture of large language models [19].

The large language models built on the Transformer architecture, which have been so successful in natural language processing [20], are readily adapted to the "language of proteins" implicit in an amino acid sequence of a protein [21]. At the heart of such protein models is an analogy to a human language: the encoding of proteins as arrangements of amino acids shares much with the way in which words are arranged into grammatically correct sentences. Both natural languages and proteins are realized as finite sequences of context-dependent “tokens” that follow a specific set of rules for their relative positions or “syntax”. The existing protein language models, trained on more than 200 million amino acid sequences, appear to successfully capture biophysical and biochemical features of proteins [22, 23] and have been applied to prediction of homology in the recently released PRotein Ortholog Search Tool (PROST) [24]. The protein secondary structure, which is determined by physicochemical properties of amino acids that dictate how neighboring amino acids interact to form secondary structure elements (alpha-helices and beta-strands), can also be presented as a protein language. In this case, the syntax is determined by how the secondary structure elements are arranged to perform the proteins’ functions [25-27]. Here, we report that the ProSSL model we developed successfully captures these “syntax” rules. By using curated datasets of two viral hallmark proteins – large terminase and portal protein – we demonstrate that ProSSL can quickly and accurately identify their homologs with low amino acid sequence identity. The predictions of the ProSSL models can be interpreted using the Shapley Additive exPlanations (SHAP) values [28, 29], which have a potential to reveal motifs and domains that are important for the homology detection task.

## Results

To build the ProSSL model, we used the standard pre-train/fine-tune schema used for large language models [30], which is outlined in **Figure 1**. Analogous to extraction of general lexical patterns, grammar, and context embodied in natural language model by pre-training on a corpus of unlabeled text (e.g., BERT [31] and RoBERTa [32] models are trained on Wikipedia), we pre-trained the ProSSL model on a database of the secondary protein structures in order to capture general patterns and relationships among secondary structure elements of proteins. Specifically, we pre-trained a Transformer-based encoder on a corpus of predicted secondary structures of the 334,881 bacterial proteins from the Swiss-Prot database [33] – a small curated database of non-redundant and representative protein sequences. The structures in the corpus were “unlabeled” – that is, the functional annotations of the proteins were removed (see **Methods** for further details). We decided to use only the encoder portion of the Transformer architecture for computational efficiency and to avoid potential overfitting issues. We tokenized the input sequences of the proteins’ secondary structure elements by “slicing” them into “meaningful chunks” (tokens, or “words”) by a tokenizer trained on the same corpus (see **Methods** for details). Following the masked language modeling (MLM) technique for training and validation [31], we replaced some randomly selected tokens in each sequence with a special “<MASK>” token, and asked the model to reconstruct the original sequence by predicting the masked tokens. During validation, the model achieved an MLM accuracy of 90.92% in exact prediction of the masked token out of the 3,000 possible tokens (in comparison, a random guess would have an accuracy of ∼0.03%).

**Figure 1.**
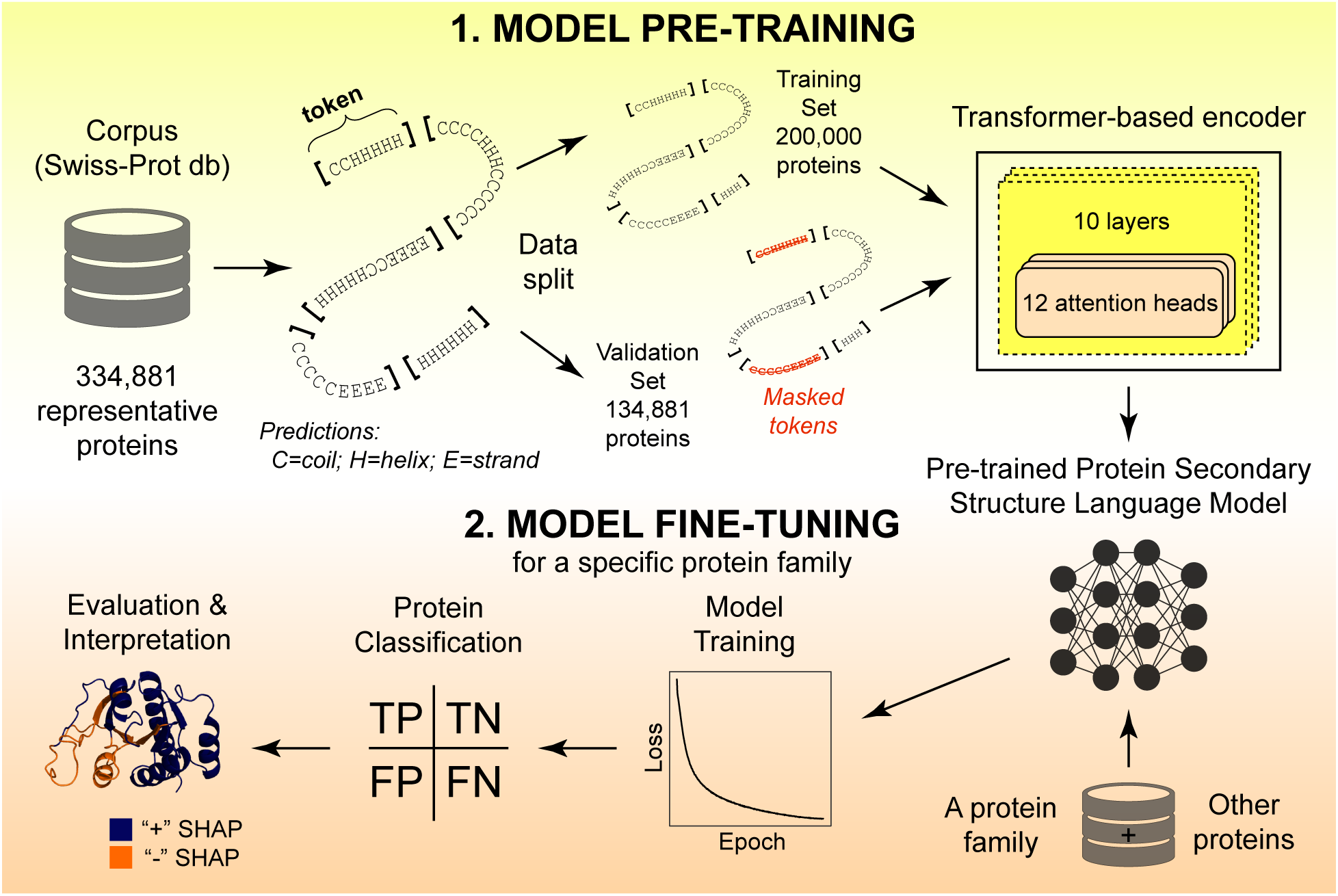
An overview of the model training, evaluation and interpretation. The training consists of two major steps. In the first step, the model was trained on tokenized secondary structures of 200,000 proteins and validated on 134, 881 proteins. In the second step, the model was fine-tuned for prediction of homologs of a protein of interest (either a terminase or a portal protein). To evaluate the model’s decision-making, SHAP values for the tokens in representative proteins were mapped to the proteins’ 3D structure.

Our pre-trained ProSSL model can be used directly for homology detection (classifying the protein into a specific protein family) without any additional training. This is achieved by setting up an “anchor” (a known member) for each protein family, and then measuring the distance between the “protein-to-classify” and each anchor to calculate the model’s confidence in the classification. Note that this so-called “few-shot” approach [34] requires knowing at least one labeled instance for each protein family. However, it is also possible to enhance the performance of the pre-trained ProSSL model in a classification task by updating the parameters of the model (fine-tuning) on a small dataset of proteins that are known to belong to a specific protein family (cf. **Figure 1** and **Methods**).

We evaluated the performance of both the pre-trained and fine-tuned ProSSL model on two classification tasks: predicting if a given protein encodes a large terminase (referred thereafter as the “terminase protein prediction” task) and predicting if a given protein encodes a portal protein (the “portal protein prediction” task). Terminase and portal proteins are viral proteins involved in the packaging of viral genome during virion assembly [35]. These proteins are found in many different viruses, and within each family, there are many examples of members that exhibit very low amino acid sequence similarity [36, 37]. We also compared the performance of the ProSSL models on these two tasks to two baselines. The first baseline is a primary-protein-structure model, which was pre-trained on the amino acid sequence representations of the proteins. This model is identical to the ProSSL model with respect to its architecture, tokenization procedure (although resulting in different “vocabularies” given the different data), hyperparameters, and training objectives. The second baseline is the aforementioned PRotein Ortholog Search Tool (PROST) [24], a primary-sequence deep learning model pre-trained on 250 million amino acid sequences that collectively contain 86 billion amino acids [23]. To our knowledge, there is currently no other secondary structure-based model to which we can compare ProSSL.

Uniform Manifold Approximation and Projection (UMAP) [38] visualization of the protein representations (embeddings) shows that both pre-trained and fine-tuned ProSSL models lead to a good separation between different types of proteins (**Figure 2** and **Figure S1**), but task-specific fine-tuning of the model results in a substantially better separation than just the pre-training of the model (**Figure S1**). Notably, the ProSSL model leads to a clearer separation between different types of proteins than our primary-protein-structure model does (**Figure 2**). These findings suggest that both fine-tuning and utilization of the secondary protein structure should result in more accurate predictions in the two classification tasks. Quantification of the model performances using Area Under the Curve (AUC) is consistent with this qualitative assessment. The fine-tuned ProSSL models have higher AUC scores than our fine-tuned primary-protein-structure models across all nine curated data splits in both tasks, and also clearly outperforms the pre-trained ProSSL model for the portal protein prediction (**Figure 3**; see **Methods**, **Table S1,** and **Table S2** for details on the construction and detailed evaluation of splits). Overall, our fine-tuned models achieve average AUC scores of 98.71-99.96% (standard deviations < 0.52%) for the terminase protein prediction and 98.62-99.98% (standard deviations < 0.67%) for the portal protein task prediction (**Figure 3**). This performance is comparable to the performance of PROST on both tasks: the average AUC scores are 99.96% vs. 99.91% for terminase protein prediction, and 99.96% vs. 99.98% for the portal protein prediction for the ProSSL and PROST models, respectively (**Figure 3**, **Table S3** and **Table S4**). It should be noted that the ProSSL model obtained comparable results with three orders of magnitude less data, making our model “lightweight” and thus easily and locally fine-tunable by a user interested to improve accuracy for homology search for a specific protein family.

**Figure 2.**
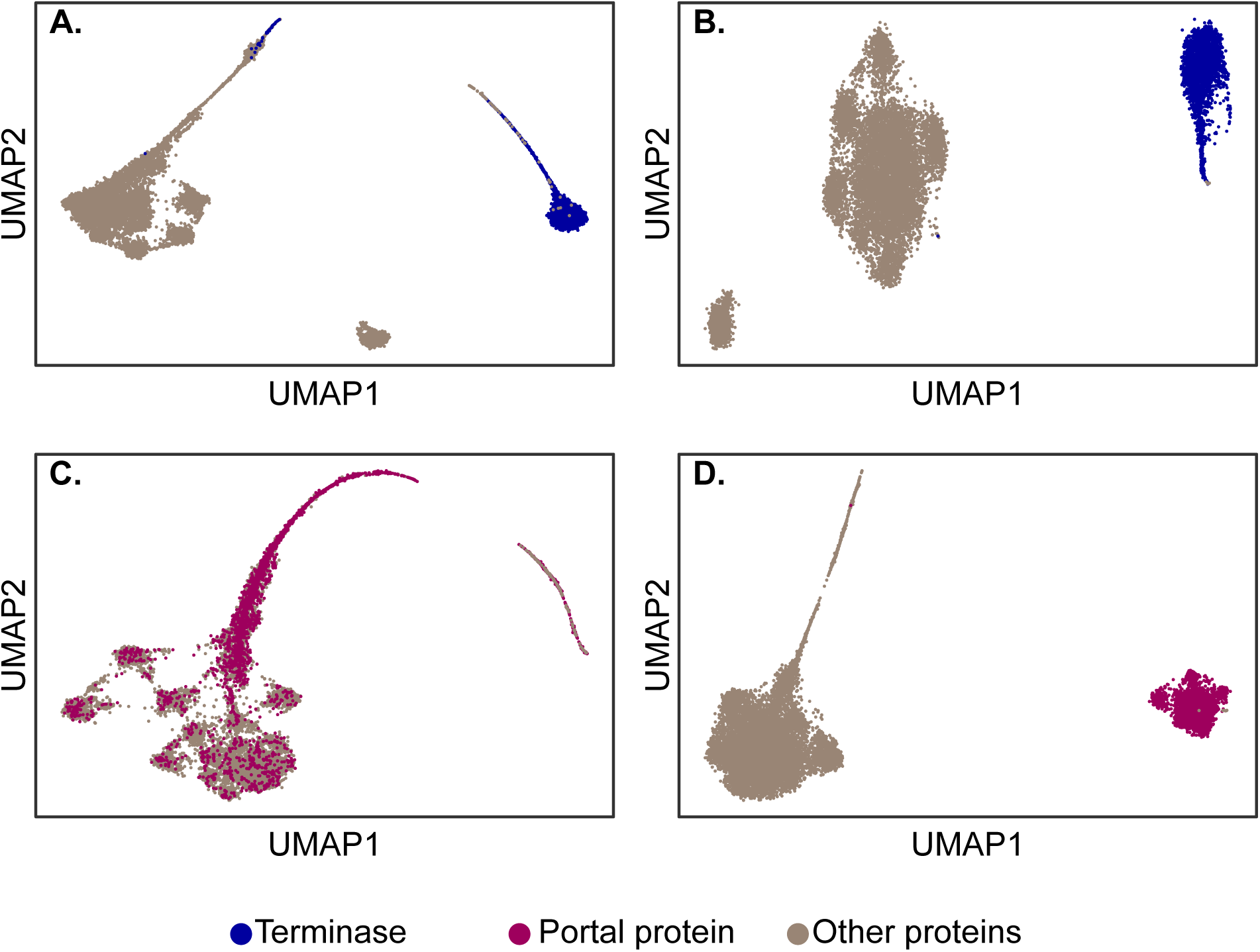
Uniform Manifold Approximation and Projection (UMAP) visualization of protein embeddings of the testing dataset for fine-tuned models. **(A)** Terminase predictions by the model trained on amino acid sequences (primary protein structure). **(B)** Terminase predictions by the model trained on the secondary protein structure. **(C)** Portal protein predictions by the model trained on the primary protein structure. **(D)** Portal protein predictions by the model trained on the secondary protein structure.

**Figure 3.**
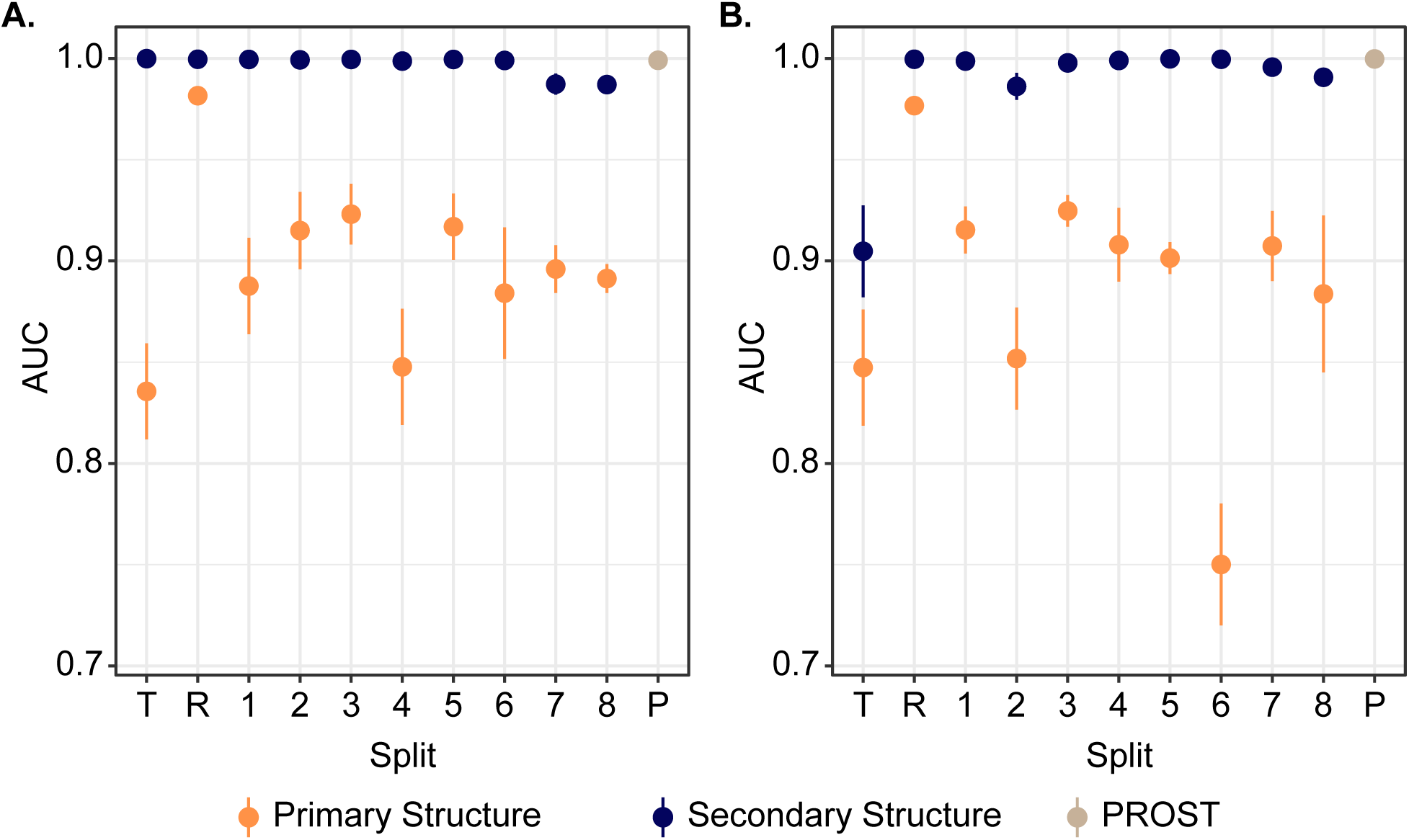
Accuracy of the models trained and tested on different subsets (splits) for (A) terminase and (B) portal proteins. On the x-axis, “T” refers to the pre-trained models, “R” refers to the models fine-tuned on the “random split”, “1”-“8” refer to the models fine-tuned on phylogenetically defined data splits 1-8, and “P” corresponds to the PROST tool. The accuracy of each model on the x-axis is measured by the AUC values (y-axis). Each dot represents the mean AUC value across replicates and error bars extend to one standard deviation from the mean. For data values, see the **Table S1 - S4**.

Collectively, the ProSSL models incorrectly classified a small number of proteins in both classification tasks (**Tables S5** and **S6**). BLASTP searches revealed that some of these false positive have significant matches to the proteins in the portal and terminase datasets due to misannotation or distant similarity. However, other false positives may represent cases of unrecognized homology. To investigate if there is a structural relatedness of the ProSSL model’s putative false positives to the known members of a protein family, we compared topological similarity of representative proteins from terminase and portal protein families to false positives, subsets of known terminases/portal proteins, and randomly selected unrelated proteins, using as a metric TM-scores of the pairwise structural alignments [14]. For both terminase and portal proteins, false positives have significantly lower TM-scores (and hence worse structural alignments) to a representative member of the protein family than known terminases/portal proteins do (**Figure 4** and **Figure 5)**. However, the false positives also tend to have higher TM-scores to a representative protein family member than the randomly sampled proteins do, and these differences are also significant for most comparisons (**Figure 4** and **Figure 5)**. This indicates the presence of some structural relationships between the reported false positives and true positive members of a protein family. To understand what those structural relationships might be, we leveraged the interpretability of the ProSSL model to examine protein features that lead to the classification of these proteins as members of either terminase or portal protein families.

**Figure 4.**
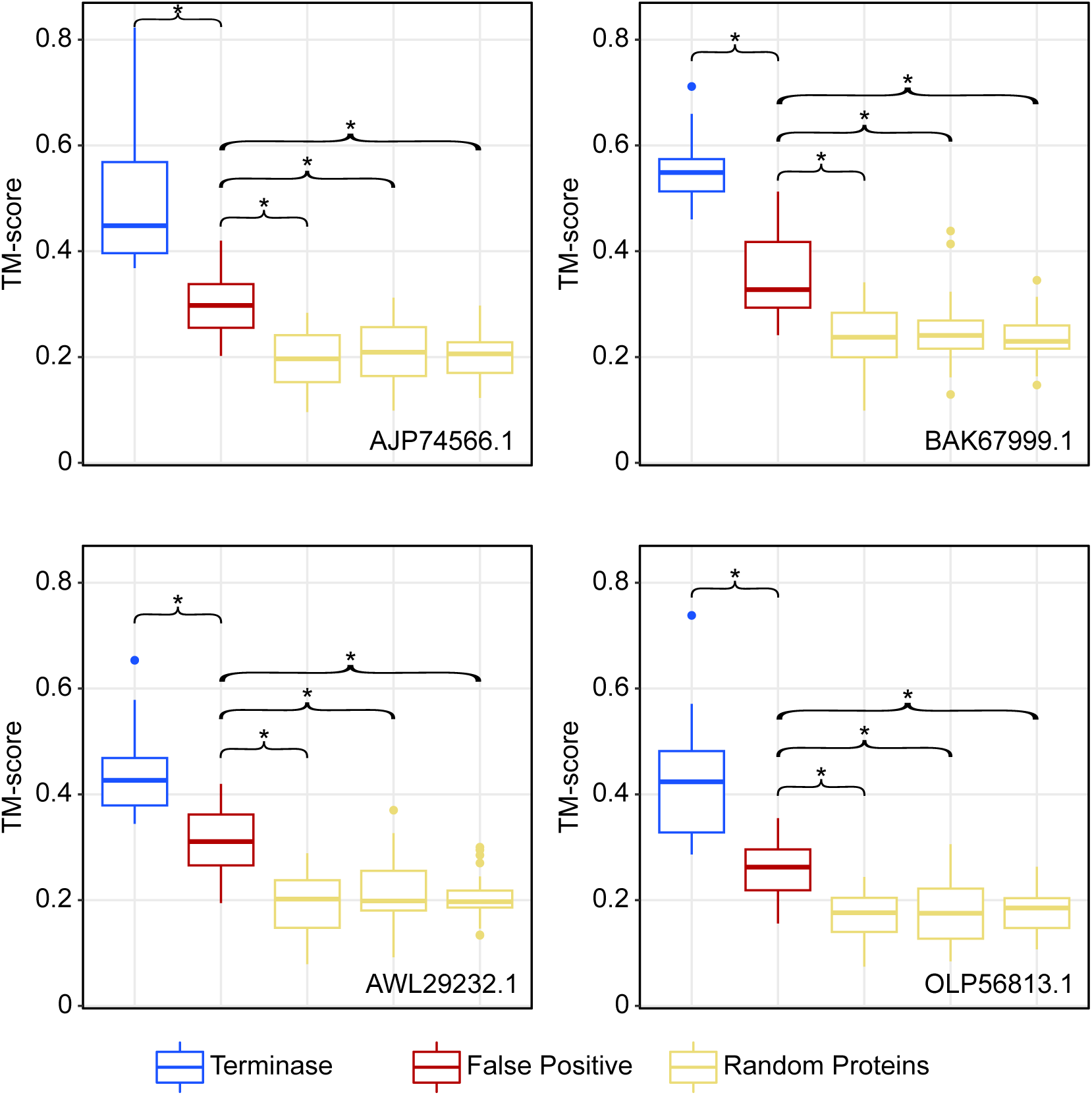
Pairwise structural similarity of four representative terminases to 25 known terminases, 23 false positives and three sets of 25 randomly selected proteins. The GenBank accessions of the representative terminases are shown in the bottom right corner of each panel. The similarity is measured via the TM-score metric. Line within a box displays the median TM-score within an analyzed dataset, and the boxes are bounded by first and third quartiles. Whiskers represent TM-scores within 1.5*interquartile range. Dots outside of whiskers are outliers. The asterisks denote the significant difference between TM-scores of two compared datasets (Mann-Whitney U-test; p-value <0.05).

**Figure 5.**
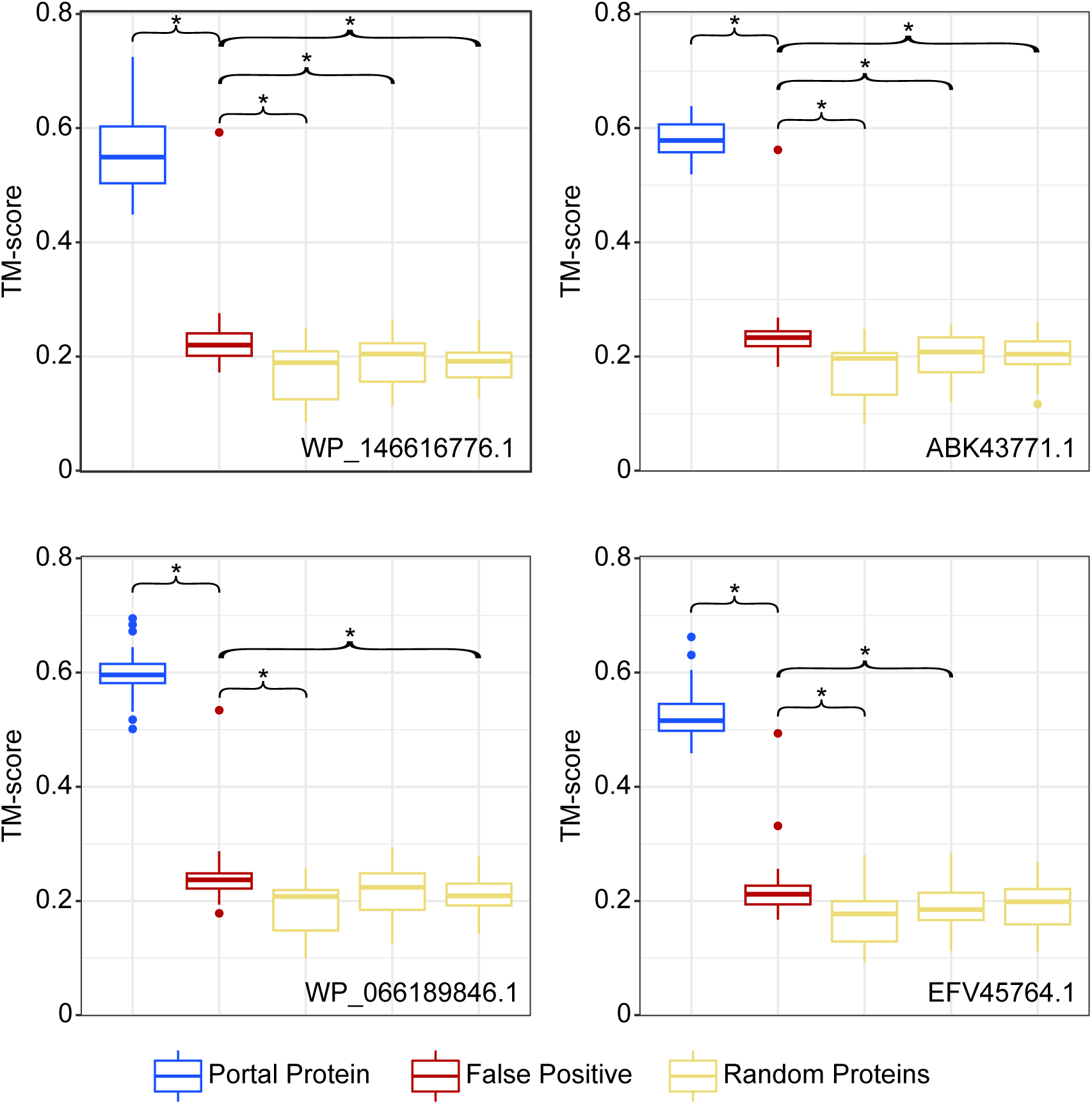
Pairwise structural similarity of four representative portal proteins to 25 known portal proteins, 60 false positives and three sets of 25 randomly selected proteins. The GenBank accessions of the representative portal proteins are shown in the bottom right corner of each panel. The similarity is measured via the TM-score metric. Line within a box displays the median TM-score within an analyzed dataset, and the boxes are bounded by first and third quartiles. Whiskers represent TM-scores within 1.5*interquartile range. Dots outside of whiskers are outliers. The asterisks denote the significant difference between TM-scores of two compared datasets (Mann-Whitney U-test; p-value <0.05).

The token representation of the proteins extracted by the ProSSL model is “interpretable” in the sense that the token encoding enables the explicit identification of protein regions responsible for the assignment of homology by the model. Such regions can be identified using the SHAP values [28] as a metric. In the context of our study, a SHAP value quantifies the individual contribution of a token to the model’s final prediction, given the entire protein sequence as input. Although SHAP values are model-specific, they can be used to gauge the influence of each segment of the protein sequence on the specific model’s ultimate decision. In the true positive matches for both the terminase and portal protein predictions, SHAP values are mostly positive throughout the protein (exemplified in **Figure 6A** and **Figure S2**), indicating that the model uses the information across the whole protein. Conversely, among the true negatives for both terminase and portal protein predictions, there is a high degree of heterogeneity in SHAP values, with many protein regions having a negative SHAP value (**Figure S3**). To understand why the model selected them in the classification tasks, we examined SHAP values of selected false positives.

**Figure 6.**
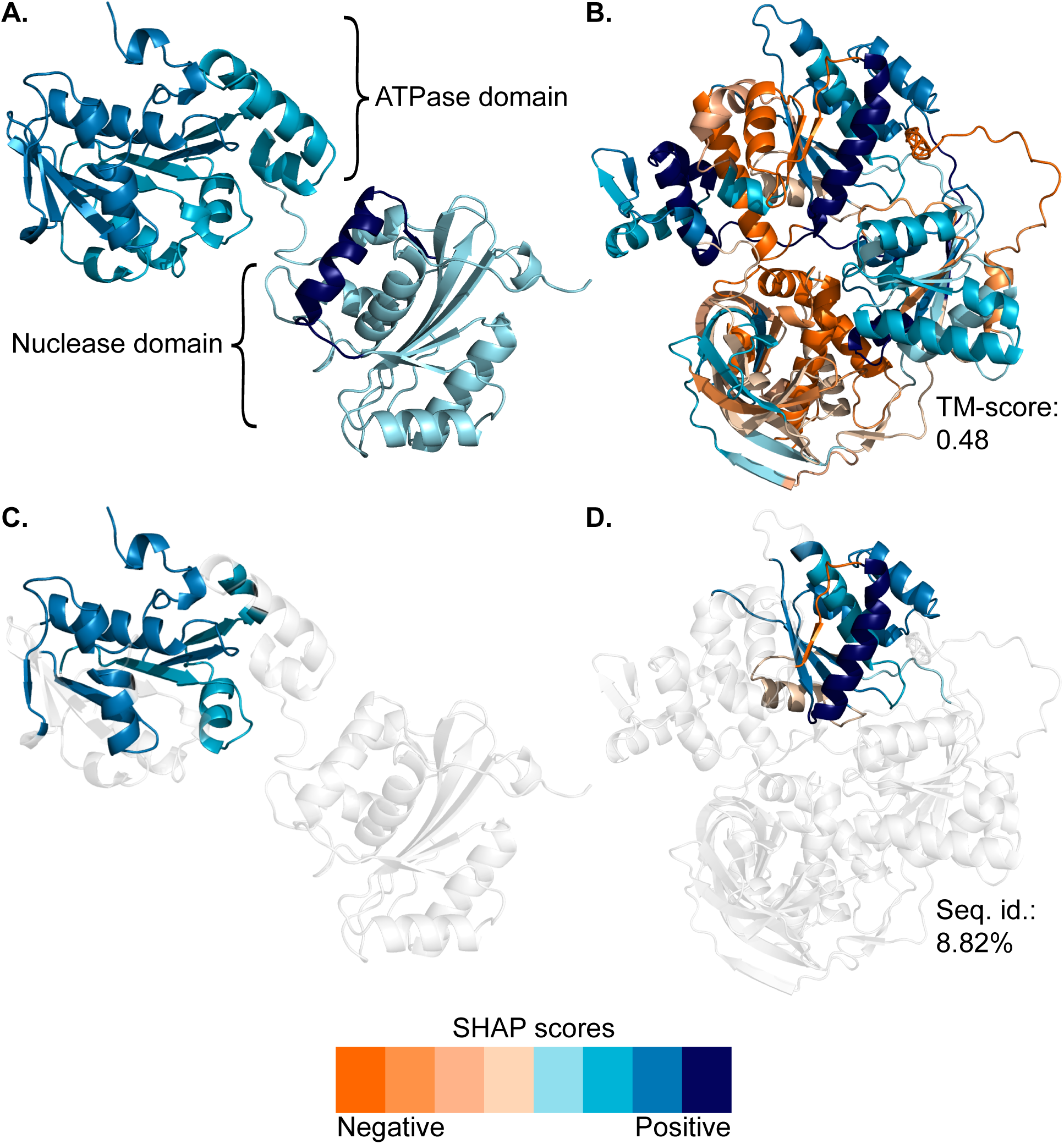
SHapley Additive exPlanations (SHAP) values of tokens of a true positive match (a representative terminase TerL) and a false positive match (the ATP-dependent RNA helicase SrmB) mapped onto their tertiary protein structures. An amino acid residue of a structure is colored according to the SHAP values of a token to which the residue belongs. The negative and positive SHAP values are shown as the gradations of orange and blue, respectively. The four gradations of orange and blue colors are assigned based on the sorting of absolute values of the SHAP values of a protein into four bins of equal size. **(A)** The structure of the true positive match (GenBank accession BAK67999.1), in which all amino acid residues are colored according to their SHAP values. **(B)** The structure of the false positive match (UniProt accession A0A399GXE0), in which all amino acid residues are colored according to their SHAP values. Its TM-score to the terminase on panel A is listed under the structure. **(C)** The structure of the true positive match (GenBank accession BAK67999.1), in which the colored segments are homologous to the regions highlighted on the structure in the panel D. **(D)** The structure of the false positive match (UniProt accession A0A399GXE0), in which the structure segments are colored only for the regions that are structurally aligned with 3 out of 4 representative terminases. “Seq. id.” refers to the sequence identity between the colored amino acids and their counterparts in the terminase on panel C. Animations of the shown structures are available as **Movies S1-S4**.

The false positive portal protein with the highest TM-scores to the representative true portal proteins (**Figure 5**) has SHAP values that are generally positive across the whole length of the protein (**Figure S4**). The protein is currently annotated as “DUF4055 domain-containing protein”, and it does not have significant amino acid sequence similarity to any of the portal proteins in our training dataset. However, in a much larger GenBank’s non-redundant (*nr*) protein database, two of its BLAST-recognizable homologs are annotated as portal proteins based on structural similarity to a known viral portal protein [39]. Hence, this “false positive” is likely a portal protein, showing that the ProSSL model can identify previously unrecognized members of a protein family.

Among 23 false positives of the terminase protein, 9 could be aligned with representative true terminases (TM-scores > 0.3). In 8 of the 9 proteins, the SHAP values from the well-aligned regions are larger than the average SHAP values of the whole protein, and in 6 of them, the difference is statistically significant (permutation test, p-value < 0.05; **Figure S5**). This indicates that structurally aligned regions generally have higher SHAP values. Moreover, in these false positives, only some segments of the protein have positive SHAP values (exemplified in **Figure 6B**). Interestingly, 8 out of the 9 false positives contain a helicase domain, and if the helicase domain is hidden by the special <MASK> token, none of them are predicted as terminases (**Table S7**). Helicases are a large group of ATP-driven enzymes that interact with the nucleic acids and catalyze various molecular processes, such as DNA unwinding and RNA metabolism [40-42]. The ATP-binding domains of both helicases and terminases belong to the large superfamily of P-loop NTPases [43] (InterPro search [44], accessed June 2023). In three closely examined alignments, the regions with the highest TM-scores are situated in the ATPase domain of both terminase and helicase (**Figures 6C, 6D,** and **S6**). While the ATP-binding domains from the P-loop NTPase superfamily are found in many protein families, our list of false positives is notable for the abundance of helicases. This is likely because both helicases and terminases belong to the Additional Strand Catalytic Glutamase (ASCE) family of the P-loop NTPase superfamily [45]. As their name implies, the NTPases from the ASCE family contain an additional beta-strand and a conserved glutamate residue [45], indicating that our model has likely learned to identify and distinguish this specific class of NTPases. Additionally, both protein classes exhibit DNA translocation activity: the ATPase domain of a terminase plays an important role in binding DNA to facilitate its translocation into a viral head [46], while helicases exhibit motor-driven translocase activity to unwind DNA molecules [47]. These functional similarities further support the likely evolutionary relatedness of these two classes of proteins. Notably, the sequence identity of the examined terminase-helicase pairs is low (8.82% for a protein shown in **Figure 6D**), and thus these pairs would not be picked up as significantly similar matches by the conventional BLAST searches [4] and HMM profiles [5]. These examples highlight the possible utility of the ProSSL model in identifying protein domains that belong to a broad class of proteins (such as a superfamily).

## Discussion

In this study, we have developed and implemented a large language model that was trained on secondary structure of proteins. While its ‘alphabet’ consists of only 3 letters, the ProSSL model is capable of learning patterns of secondary structure arrangements in bacterial proteins. We showed that even when trained on a small number of proteins, the model successfully detects homologs, including very distant and previously unrecognized ones. Notably, the ProSSL model outperforms the primary protein structure model on the comparable datasets. Taken together, these observations suggest that the ProSSL model could provide an excellent complement to the amino acid sequence models. A combined primary and secondary protein structure models may further increase sensitivity of homology searches. Moreover, the ProSSL model can allow more sensitive, structure-based predictions for classes of proteins whose 3D structures are not yet available due to technical challenges.

We also showed that the relevant protein secondary structure features of a match can be spatially localized to a specific region of a protein (for example, a domain or a fold) by mapping the SHAP values onto amino acid sequences or 3D protein structures (if available). While SHAP values are model-specific, they can provide a researcher with a measure of confidence about the quality and specific regions of similarity of the returned matches under the utilized model. In the future, model-agnostic interpretation metrics need to be developed to allow comparison of matches across models trained on different datasets.

As we demonstrated for the case of distant homology between terminases and helicases, the ProSSL model can pick up similarity of specific domains or folds within a protein. Training and fine-tuning the model on domains rather than full-length proteins will expand its applicability to eukaryotic proteins, many of which consist of multiple domains.

Finally, while we illustrated the utility of the ProSSL model on the task of homology detection, the model is not limited to just that task. For example, search for signal peptides was shown to be improved by application of the protein language models [48] and is likely to benefit from the secondary structure information due to its importance for the signal sequence functionality [49, 50].

## Methods

### Protein datasets for model training and evaluation

For the pre-training of the ProSSL and primary protein structure models, 335,576 amino acid sequences of bacterial proteins were retrieved from the Swiss-Prot database (accessed June 2022) [33, 51]. Sequences that contained ambiguous amino acids or had a length of either above 5,000 or below 50 amino acids were removed, resulting in 334,881 amino acid sequences. Of these, 200,000 sequences were randomly selected (the random seed of 42) for a training dataset, while the remaining 134,881 sequences were used for validation.

For fine-tuning of the pre-trained models for the “terminase protein prediction” task, the curated dataset of 11,051 amino acid sequences of large terminases (TerL) (referred throughout the manuscript as “terminases” for brevity) was retrieved from ref. [37]. Of these, 11,002 sequences without any ambiguous amino acids were retained. To create a ‘non-terminase’ dataset, 50,055 proteins that were not annotated as a ‘terminase’ were retrieved from the UniRef50 database [52] (accessed April 2023). Of these, 49,836 sequences with lengths ranging between 50 and 5,000 amino acids and without ambiguous amino acids were retained.

For fine-tuning of the pre-trained models for the “portal protein prediction” task, the curated dataset of 14,745 amino acid sequences of portal proteins was retrieved from ref. [53]. Of these, 14,697 sequences without any ambiguous amino acids were retained. To construct the ‘non-portal’ dataset, nine sequences that were annotated as ‘portal protein’ were removed from the above-described ‘non-terminase’ dataset, resulting in 49,827 protein sequences.

To increase the genetic distance between the training and testing datasets in the fine-tuning for protein prediction tasks, eight data splits were designed using the phylogenies of terminases and portal proteins. The phylogenies were obtained from the refs. [37] and [53]. For splits 1, 3, 5, and 7, proteins from a specific subtree were used for the testing, while proteins from the rest of the tree were used for the training (**Figures S7** and **S8**). For splits 2, 4, 6, and 8, proteins branching between subtrees that contain testing and training data were completely removed (**Figures S7** and **S8**). In addition to these manually curated splits, we also created a “random split” of the data by randomly dividing the terminase and portal protein datasets into training and test sets, comprising 80% and 20% of the data, respectively (again, using a random seed of 42).

For the primary protein structure models, the amino acid sequences were used as is. For the ProSSL models, the secondary structures of the proteins in the above-described datasets were predicted using Porter5 [17]. The secondary structure of each protein was encoded as a string of text, in which each amino acid was replaced with a single letter representing the type of secondary structure element the amino acid is predicted to be in (“H” for helix, “E” for beta-strand and “C” for coil).

### Tokenization

Unlike the situation with words in a natural language, for proteins, it is not *a priori* clear as to what might constitute an appropriate token. The process of defining the vocabulary for our model involved segmenting a training set of 200,000 proteins into shorter, contiguous character sequences. The most recurrent sequences were then incorporated into the vocabulary. To determine the optimal vocabulary size, we conducted ten experimental trials, exploring sizes from 1,000 to 10,000, increasing in increments of 1,000. The tokenization was carried out using SentencePiece [54], a language-agnostic tokenizer commonly used for natural language processing that works on raw input text and does not require pre-tokenization. The suitability of the vocabulary size was evaluated using entropy, which measures the minimum theoretical number of bits required per symbol to encode information from a given data source [55]. The tokenizers were subsequently validated on the validation set of 134,881 proteins.

For the protein secondary structure data, using 1,000 tokens resulted in a high tokenizer entropy of over 50. At 3,000 tokens, the entropy dropped to 9.36 and remained roughly unchanged regardless of further increases in the vocabulary size. For the protein primary structure data, at the vocabulary size of 5,000 the entropy dropped to 10.30 and remained approximately unchanged at larger vocabulary sizes (**Figure S9**). These entropy values are similar to the entropy of 9.12 when the tokenizer was evaluated on a standard text corpus of Wikipedia articles [56]. Therefore, vocabulary sizes of 3,000 and 5,000 were selected for the secondary and the primary structure data, respectively.

### Model Architecture

For computational efficiency and to avoid potential overfitting issues, only the encoder portion of the Transformer architecture was used. Specifically, a structure consisting of 10 layers, each featuring 12 attention heads was employed, mirroring the architecture of the widely recognized RoBERTa-base language model [32]. The model’s hidden dimension was configured to be 768. Each of the 12 attention heads in a layer parsed the input (tokenized sequences of either amino acids or protein secondary structure elements) and extracted the most informative and important features of each token in each sequence. The results of all attention heads in a layer were consolidated and passed to the next layer in the model. After the process was repeated across the 10 layers, the model summarized the progressively refined and abstracted features of each sequence in a high-dimensional vector (“latent representation”). These vectors were the output of the model. All models were coded in Python using the Hugging Face’s Transformers library [57].

### Pre-Training Procedure

To train our model, the masked language modeling (MLM) method [31] with random initialization (i.e., without providing any prior knowledge about the input sequences) was used. Specifically, in each of the tokenized input sequences, 15% of randomly selected tokens (using a random seed of 42) were replaced with a special "<MASK>" token. The models were then trained to predict the masked tokens using the context provided by the other unmasked tokens. A learning rate of 1e-5 and a batch size of 128 were used, while other hyperparameters were set to the RoBERTa models’ defaults, as specified in ref. [32]. The training was carried out on the 200,000 proteins of the training set. After the training, the models were evaluated using a masked-token reconstruction task on 134,881 proteins of the validation set. The performance was assessed by calculating MLM accuracy [31], which measures how well the models could predict masked tokens in the unseen protein sequences.

### Fine-Tuning of Models for Protein Classification Tasks

The pre-trained models were fine-tuned for the tasks of predicting proteins as either terminases or portal proteins. To do so, a specialized output layer (a prediction head) was added to the pre-trained model. The prediction head and the encoder were then jointly fine-tuned on the above-described training data. A learning rate of 5e-5 and batch size of 16 were used, with the training process lasting for 10 epochs. An early stopping was used to halt the training if the performance on a validation dataset stopped improving.

To ensure that performances of our models were not due to any specific split of the data, the models were tested using eight different curated data splits and one randomly split data set for both the terminase and portal protein identification tasks (the construction of the splits is described in the **“Protein datasets for model training and evaluation”** section of the Methods). Each experiment was repeated five times with different random seeds to ensure the robustness and repeatability of our evaluations.

The effectiveness of the models was evaluated using AUC (Area Under the Curve) and Uniform Manifold Approximation and Projection (UMAP) [38].

### Comparison to PROST

The PROST program [24] was downloaded in May 2023. For each of 10 randomly selected terminase proteins, local PROST searches against databases constructed from all proteins in “terminase” and “non-terminase” datasets were conducted. The results of both searches were combined and sorted by the PROST distance between the query and the subjects. AUC value was calculated from these sorted distances (**Table S3**). The 10 randomly selected portal proteins were analyzed using the same procedure and databases constructed from all proteins in “portal” and “non-portal” datasets (**Table S4**).

### Interpretation of Model Predictions using Shapley Values

The SHAP values for each input token in a protein of interest were calculated using the SHAP (Shapley Additive exPlanations) framework implemented in the SHAP package (version 0.41.0) [28], with all adjustable hyper-parameters set to the default settings. SHAP values were derived through a series of computational experiments, in which different portions of the input sequence were ablated. Tokens with significant impact on the model’s prediction were assigned high absolute SHAP values, while those with minimal influence received low absolute SHAP values.

### Structural Comparison of False Positives to representative Terminase and Portal Proteins

False positives of terminase and portal proteins pooled from the results of all fine-tuned ProSSL models were used as queries in a BLASTP v2.5 search [4] (e-value < 0.1) against the terminase and portal protein datasets, respectively; only false positives without any significant matches were retained (**Table S5** and **S6**).

The false positives were compared to four known terminases and four known portal proteins, which were randomly selected from four distinct phylogenetically informed data splits (specifically, splits 1, 3, 5 and 7) to ensure a diverse representation of these protein families. These proteins are referred to as “representative proteins”. Pairwise structural alignments between each representative protein and each false positive were calculated using TM-align [14]. The TM-scores were normalized by the length of the representative protein. False positives with the TM-score larger than 0.3 in at least 3 out of 4 alignments with the representative proteins were retained for further analyses. Only regions of a false positive that were aligned with at least 3 out of 4 representative proteins and consisted of more than 30 consecutive amino acids were considered to be “structurally aligned”.

Additionally, TM-scores were calculated between the representative proteins and 25 terminases and 25 portal proteins with available tertiary structures that were randomly sampled from terminase and portal datasets, as well as between the representative proteins and 3 sets of 25 proteins randomly sampled without replacement from the “non-portal” dataset.

Tertiary structures of all selected proteins were downloaded from the AlphaFold database (accessed June 2023) [10]. Visualization of protein structures and mapping of SHAP values onto them was carried out using PyMOL v2.5 [58].

For one false positive portal protein (UniProt accession A0A0Q4N4T9), a BLASTP [4] search of the *nr* database with the default settings was carried out using the NCBI web page (https://blast.ncbi.nlm.nih.gov; accessed July 2023).

## Data and Code Availability

Amino acid sequences used to train and evaluate models, the trained ProSSL model, Python code for fine-tuning of the model and Shapley value calculations, as well as all Python scripts needed to reproduce presented analyses of the two viral protein families, are publicly available in a GitHub repository at https://github.com/soroushv-dartmouth/LanguageOfProteins.

## Acknowledgements

We thank Jonathan Chiou (Dartmouth class of 2022) for his initial exploration of the application of deep learning techniques to homology prediction as part of his senior undergraduate thesis research, which led to initialization of this project.

## Supplementary Figure Legends

**Figure S1. Uniform Manifold Approximation and Projection (UMAP) visualization of protein embeddings of the testing dataset for pre-trained models. (A)** Terminase predictions by the model pre-trained on amino acid sequences (primary protein structure). **(B)** Terminase predictions by the model pre-trained on the secondary protein structure. **(C)** Portal protein predictions by the model pre-trained on the primary protein structure. **(D)** Portal protein predictions by the model pre-trained on the secondary protein structure.

**Figure S2. SHAP values of the tokens for a true positive portal protein match (GenBank accession ABK43771.1) mapped onto the protein’s tertiary structure.** An amino acid residue of a structure is colored according to the SHAP value of a token to which the residue belongs. The negative and positive SHAP values are shown in the gradations of orange and blue, respectively. The four gradations of orange and blue colors are assigned to the residue based on the sorting of absolute values of this protein’s SHAP values into four bins of equal size.

**Figure S3. SHAP values of the tokens for representative true negative matches from the terminase and portal protein classification tasks mapped onto their tertiary protein structures.** An amino acid residue of a structure is colored according to the SHAP value of a token to which the residue belongs. The negative and positive SHAP values are shown in the gradations of orange and blue, respectively. The four gradations of orange and blue colors are assigned to the residue based on the sorting of absolute values of the SHAP values of a protein into four bins of equal size. **(A)** The structure of the true negative match (UniProt accession I1CZL4) from the terminase classification task, in which all amino acid residues are colored according to their SHAP values. **(B)** The structure of the true negative match (UniProt accession A0A376AF53) from the portal protein classification task, in which all amino acid residues are colored according to their SHAP values.

**Figure S4. SHAP values of the tokens for the false positive match in the portal protein classification tasks with the highest TM-score (UniProt accession A0A0Q4N4T9) mapped on its tertiary protein structure.** An amino acid residue of a structure is colored according to the SHAP values of a token to which the residue belongs. The negative and positive SHAP values are shown in the gradations of orange and blue, respectively. The four gradations of orange and blue colors are assigned to the residue based on the sorting of absolute values of the SHAP values of the protein into four bins of equal size. **(A)** The SHAP values from the model trained on the phylogenetically informed split #1. **(B)** The SHAP values from the model trained on the phylogenetically informed split #2.

**Figure S5. Evaluation of SHAP values in within regions that structurally align with at least 3 out of 4 representative terminases.** Each panel represent one of the nine evaluated false positives, selected as having a TM-score >0.3 with at least 3 of 4 representative terminases. The UniProt accessions of the false positives are shown on top of each graph. For each false positive, the SHAP value for the aligned regions (normalized for its length) is shown as dashed blue line and the SHAP value of the whole protein (also normalized for its length) is shown in dashed red line. The SHAP value per residue in 1,000 random shuffles of amino acid residues (and their SHAP values) are shown as a distribution in brown color. P-values designate the significance of the difference between the true and shuffled SHAP values under the permutation tests.

**Figure S6. SHAP values of the tokens for a true positive match (a representative terminase) and two false positive matches (helicases) mapped onto their tertiary protein structures.** An amino acid residue of a structure is colored according to the SHAP value of a token to which the residue belongs. The negative and positive SHAP values are shown in the gradations of orange and blue, respectively. The four gradations of orange and blue colors are assigned to the residue based on the sorting of absolute values of the SHAP values of a protein into four bins of equal size. **(A)** The structure of the true positive match (GenBank accession BAK67999.1), in which the colored segments are homologous to the regions highlighted on the structure in the panel B. **(B)** The structure of the false positive match (UniProt accession A0A7K0CVX0), in which the structure segments are colored only for the regions that are structurally aligned with 3 out of 4 representative terminases. **(C)** The structure of the true positive match (GenBank accession BAK67999.1), in which the colored segments are homologous to the regions highlighted on the structure in the panel D. **(D)** The structure of the false positive match (UniProt accession A4FJZ7), in which the structure segments are colored only for the regions that are structurally aligned with 3 out of 4 representative terminases.

**Figure S7. Phylogenetically informed design of the data splits 1-8 of the terminase dataset used for the training and testing of the models.** The phylogenetic tree relates all used terminases and is the same on all panels. For each data split, black branches show terminases used for the model training, red branches correspond to terminases used for the model testing, and gray branches refer to terminases that were not used neither in the model testing nor training. The phylogenetic tree is scaled with number of substitutions per site (see scale bar) and was rooted using the testing set of the split 1.

**Figure S8. Phylogenetically informed design of the data splits 1-8 of the portal protein dataset used for the training and testing of the models.** The phylogenetic tree relates all used portal proteins and is the same on all panels. For each data split, black branches show portal proteins used for the model training, red branches correspond to portal proteins used for the model testing, and gray branches refer to portal proteins that were not used neither in the model testing nor training. The phylogenetic tree is scaled with number of substitutions per site (see scale bar) and was rooted using the testing set of the split 1.

**Figure S9. Entropy values of models pre-trained on different vocabulary sizes.** The points correspond to the tested vocabulary sizes, which ranged from 1,000 to 10,000 tokens.

## Supplementary Table Captions

**Table S1. Evaluation of accuracy of the "terminase protein prediction" task across nine splits and under two fine-tuned large language models.**

**Table S2. Evaluation of accuracy of the "portal protein prediction" task across nine splits and under two fine-tuned large language models.**

**Table S3. AUC values for 10 randomly selected terminase proteins, evaluated under three models.**

**Table S4. AUC values for 10 randomly selected portal proteins, evaluated under three models.**

**Table S5. All false positives from the "terminase protein prediction" task carried out under the ProSSL model.**

**Table S6. All false positives from the "portal protein prediction" task carried out under the ProSSL model.**

**Table S7. Eight false positive terminases with the helicase domain, classified as "non-terminase" after its masking with the special <MASKED> token.**

## Notes

https://github.com/soroushv-dartmouth/LanguageOfProteins

### Competing Interest Statement

The authors have declared no competing interest.

