## Supplementary Figures S1-S9 for "Homology detection using a protein secondary structure-based large language model"

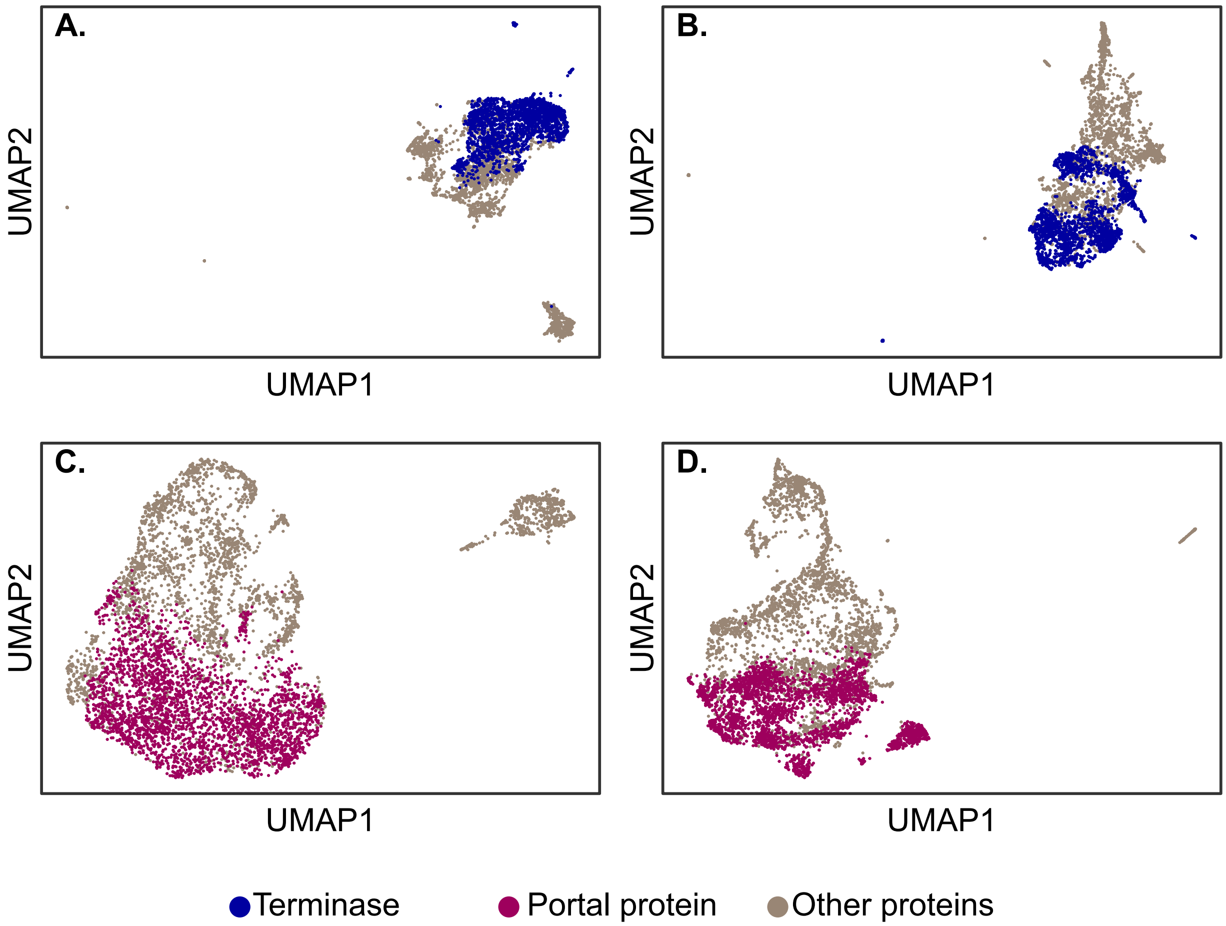


**Figure S1**. **Uniform Manifold Approximation and Projection (UMAP) visualization of protein embeddings of the testing dataset for pre-trained models. (A)** Terminase predictions by the model pre-trained on amino acid sequences (primary protein structure). **(B)** Terminase predictions by the model pre-trained on the secondary protein structure. **(C)** Portal protein predictions by the model pre-trained on the primary protein structure. **(D)** Portal protein predictions by the model pre-trained on the secondary protein structure.

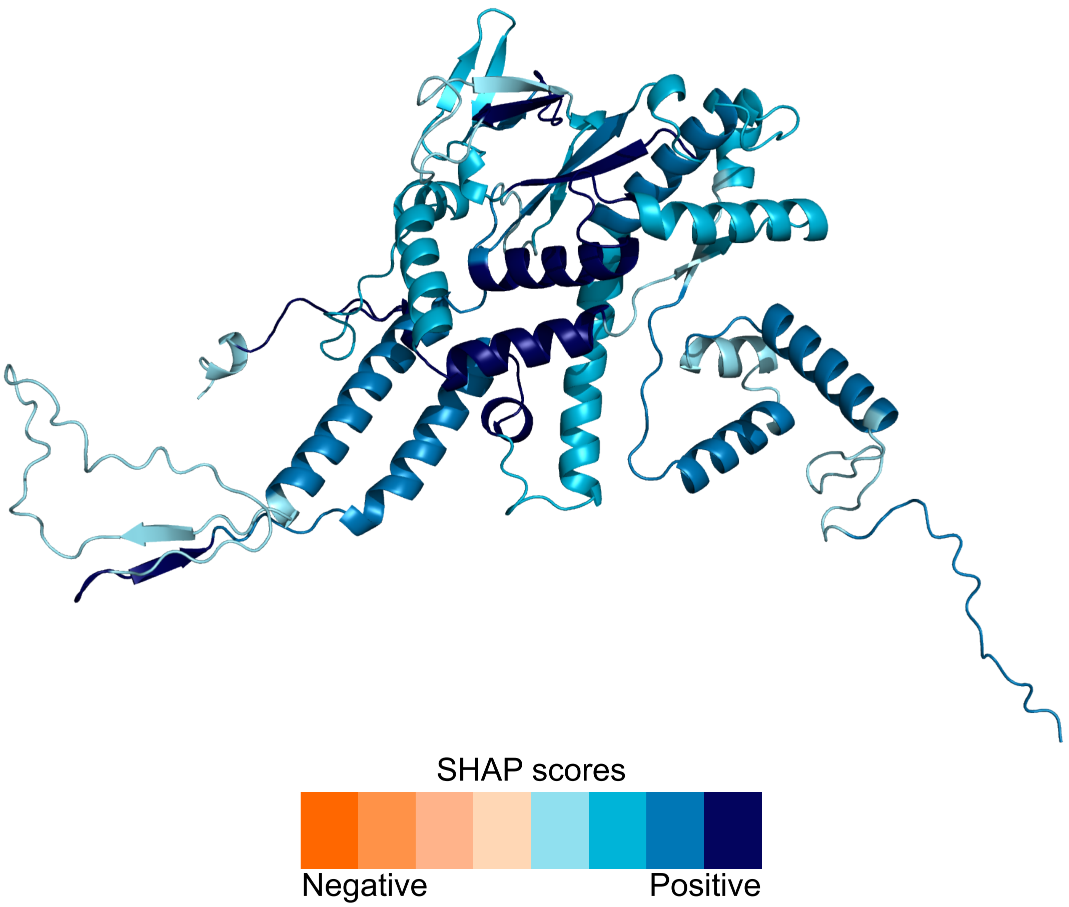


**Figure S2. SHAP values of the tokens for a true positive portal protein match (GenBank accession ABK43771.1) mapped onto the protein's tertiary structure.** An amino acid residue of a structure is colored according to the SHAP value of a token to which the residue belongs. The negative and positive SHAP values are shown in the gradations of orange and blue, respectively. The four gradations of orange and blue colors are assigned to the residue based on the sorting of absolute values of this protein's SHAP values into four bins of equal size.

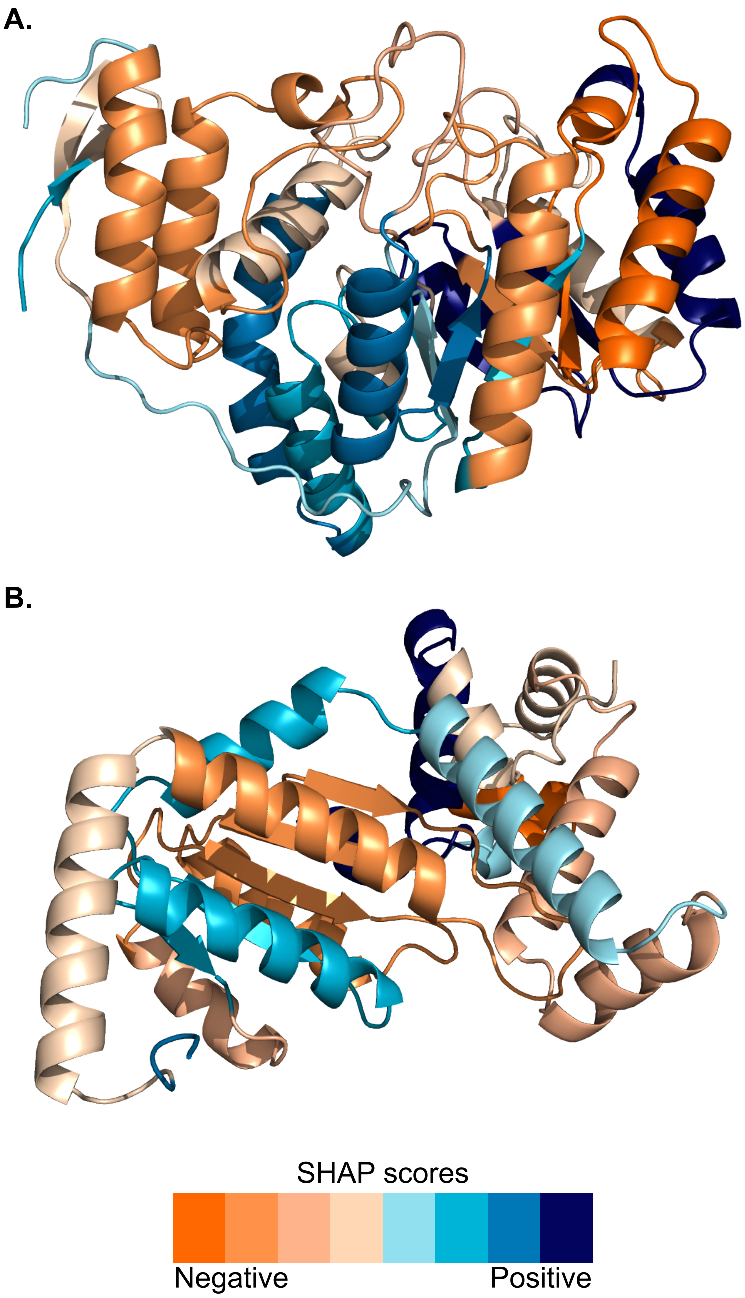


**Figure S3. SHAP values of the tokens for representative true negative matches from the terminase and portal protein classification tasks mapped onto their tertiary protein structures.** An amino acid residue of a structure is colored according to the SHAP value of a token to which the residue belongs. The negative and positive SHAP values are shown in the gradations of orange and blue, respectively. The four gradations of orange and blue colors are assigned to the residue based on the sorting of absolute values of the SHAP values of a protein into four bins of equal size. **(A)** The structure of the true negative match (UniProt accession I1CZL4) from the terminase classification task, in which all amino acid residues are colored according to their SHAP values. **(B)** The structure of the true negative match (UniProt accession A0A376AF53) from the portal protein classification task, in which all amino acid residues are colored according to their SHAP values.

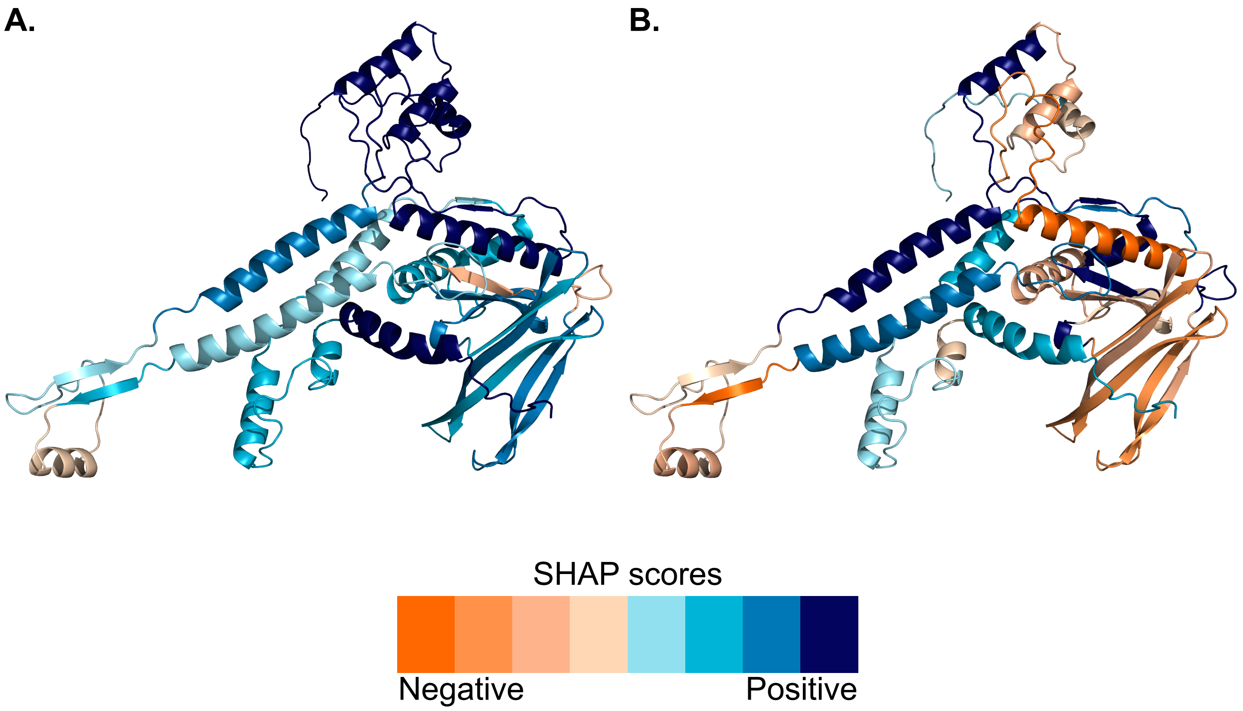


**Figure S4. SHAP values of the tokens for the false positive match in the portal protein classification tasks with the highest TM-score (UniProt accession A0A0Q4N4T9) mapped on its tertiary protein structure.** An amino acid residue of a structure is colored according to the SHAP values of a token to which the residue belongs. The negative and positive SHAP values are shown in the gradations of orange and blue, respectively. The four gradations of orange and blue colors are assigned to the residue based on the sorting of absolute values of the SHAP values of the protein into four bins of equal size. **(A)** The SHAP values from the model trained on the phylogenetically informed split #1. **(B)** The SHAP values from the model trained on the phylogenetically informed split #2.

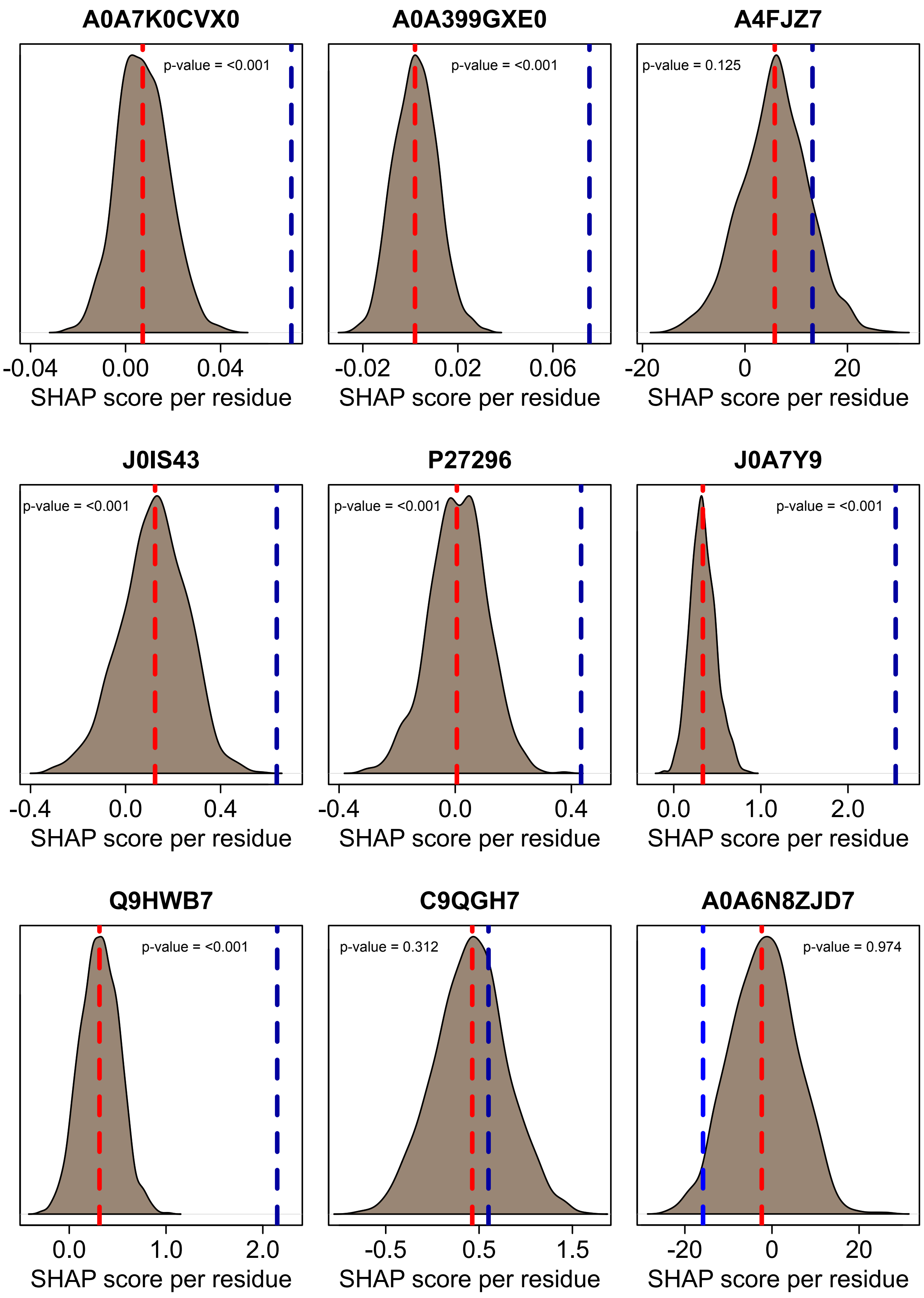
A0A7K0CVX0
p-value = <0.001
-0.04
0.00
0.04
SHAP score per residue
A0A399GXE0
p-value = <0.001
-0.02
0.02
0.06
SHAP score per residue
A4FJZ7
p-value = 0.125
-20
0
20
SHAP score per residue
J0IS43
p-value = <0.001
-0.4
0.0
0.4
SHAP score per residue
P27296
p-value = <0.001
-0.4
0.0
0.4
SHAP score per residue
J0A7Y9
p-value = <0.001
0.0
1.0
2.0
SHAP score per residue
Q9HWB7
p-value = <0.001
0.0
1.0
2.0
SHAP score per residue
C9QGH7
p-value = 0.312
-0.5
0.5
1.5
SHAP score per residue
A0A6N8ZJD7
p-value = 0.974
-20
0
20
SHAP score per residue

**Figure S5. Evaluation of SHAP values in within regions that structurally align with at least 3 out of 4 representative terminases.** Each panel represent one of the nine evaluated false positives, selected as having a TM-score >0.3 with at least 3 of 4 representative terminases. The UniProt accessions of the false positives are shown on top of each graph. For each false positive, the SHAP value for the aligned regions (normalized for its length) is shown as dashed blue line and the SHAP value of the whole protein (also normalized for its length) is shown in dashed red line. The SHAP value per residue in 1,000 random shuffles of amino acid residues (and their SHAP values) are shown as a distribution in brown color. P-values designate the significance of the difference between the true and shuffled SHAP values under the permutation tests.

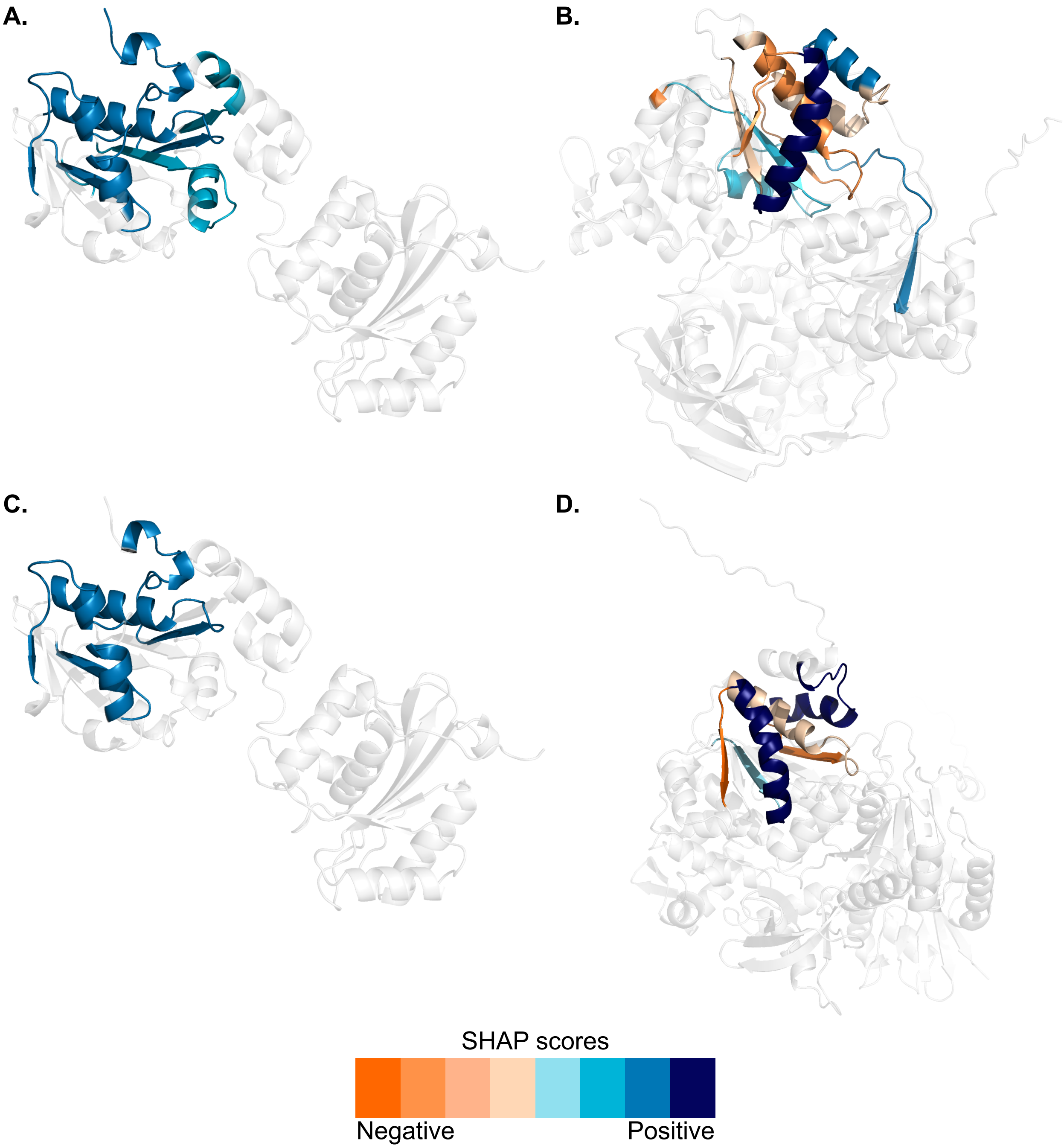


**Figure S6. SHAP values of the tokens for a true positive match (a representative terminase) and two false positive matches (helicases) mapped onto their tertiary protein structures.** An amino acid residue of a structure is colored according to the SHAP value of a token to which the residue belongs. The negative and positive SHAP values are shown in the gradations of orange and blue, respectively. The four gradations of orange and blue colors are assigned to the residue based on the sorting of absolute values of the SHAP values of a protein into four bins of equal size. **(A)** The structure of the true positive match (GenBank accession BAK67999.1), in which the colored segments are homologous to the regions highlighted on the structure in the panel B. **(B)** The structure of the false positive match (UniProt accession A0A7K0CVX0), in which the structure segments are colored only for the regions that are structurally aligned with 3 out of 4 representative terminases. **(C)** The structure of the true positive match (GenBank accession BAK67999.1), in which the colored segments are homologous to the regions highlighted on the structure in the panel D. **(D)** The structure of the false positive match (UniProt accession A4FJZ7), in which the structure segments are colored only for the regions that are structurally aligned with 3 out of 4 representative terminases.

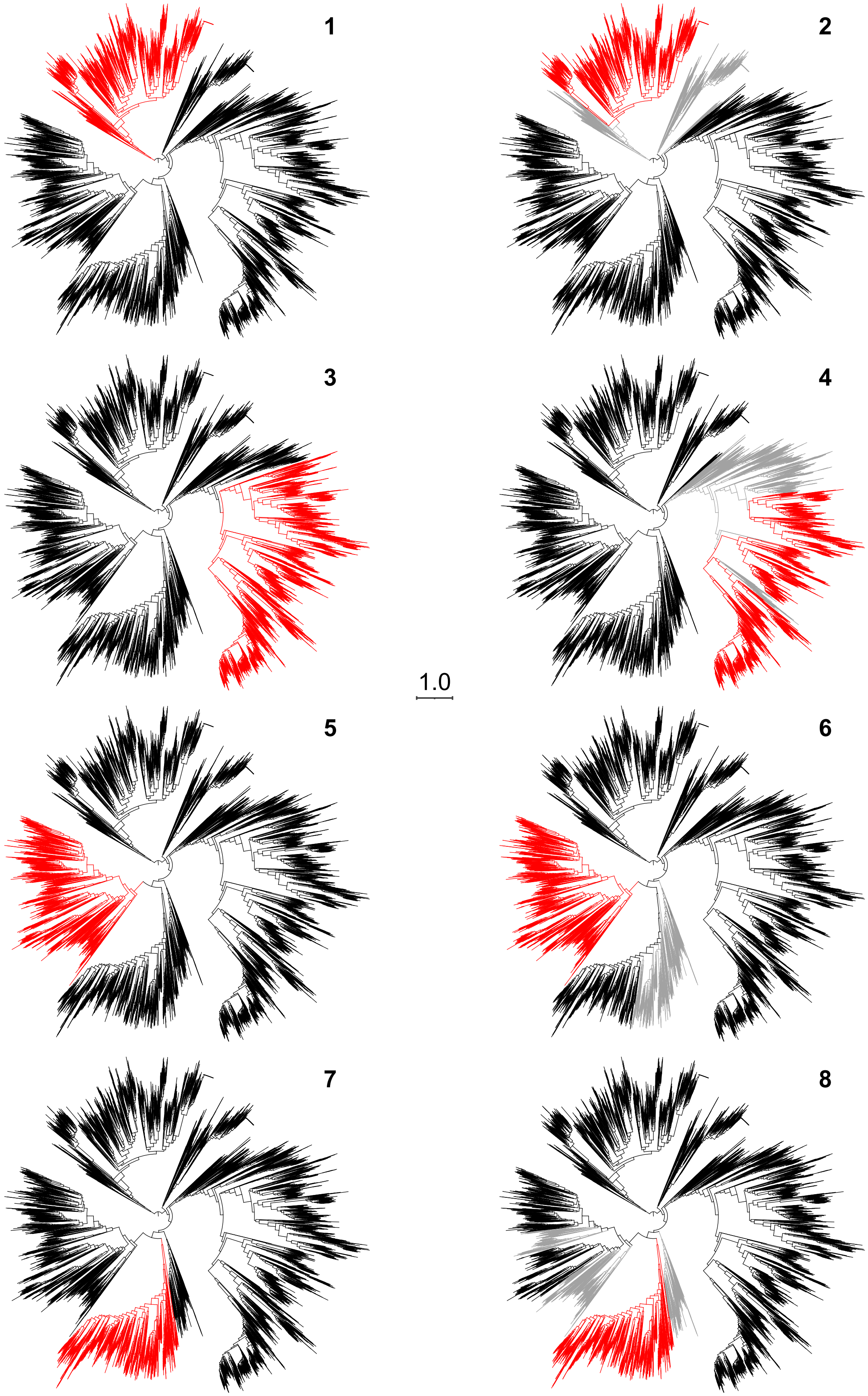
1
2
3
4
1.0
5
6
7
8

**Figure S7**. **Phylogenetically informed design of the data splits 1-8 of the terminase dataset used for the training and testing of the models.** The phylogenetic tree relates all used terminases and is the same on all panels. For each data split, black branches show terminases used for the model training, red branches correspond to terminases used for the model testing, and gray branches refer to terminases that were not used neither in the model testing nor training. The phylogenetic tree is scaled with number of substitutions per site (see scale bar) and was rooted using the testing set of the split 1.

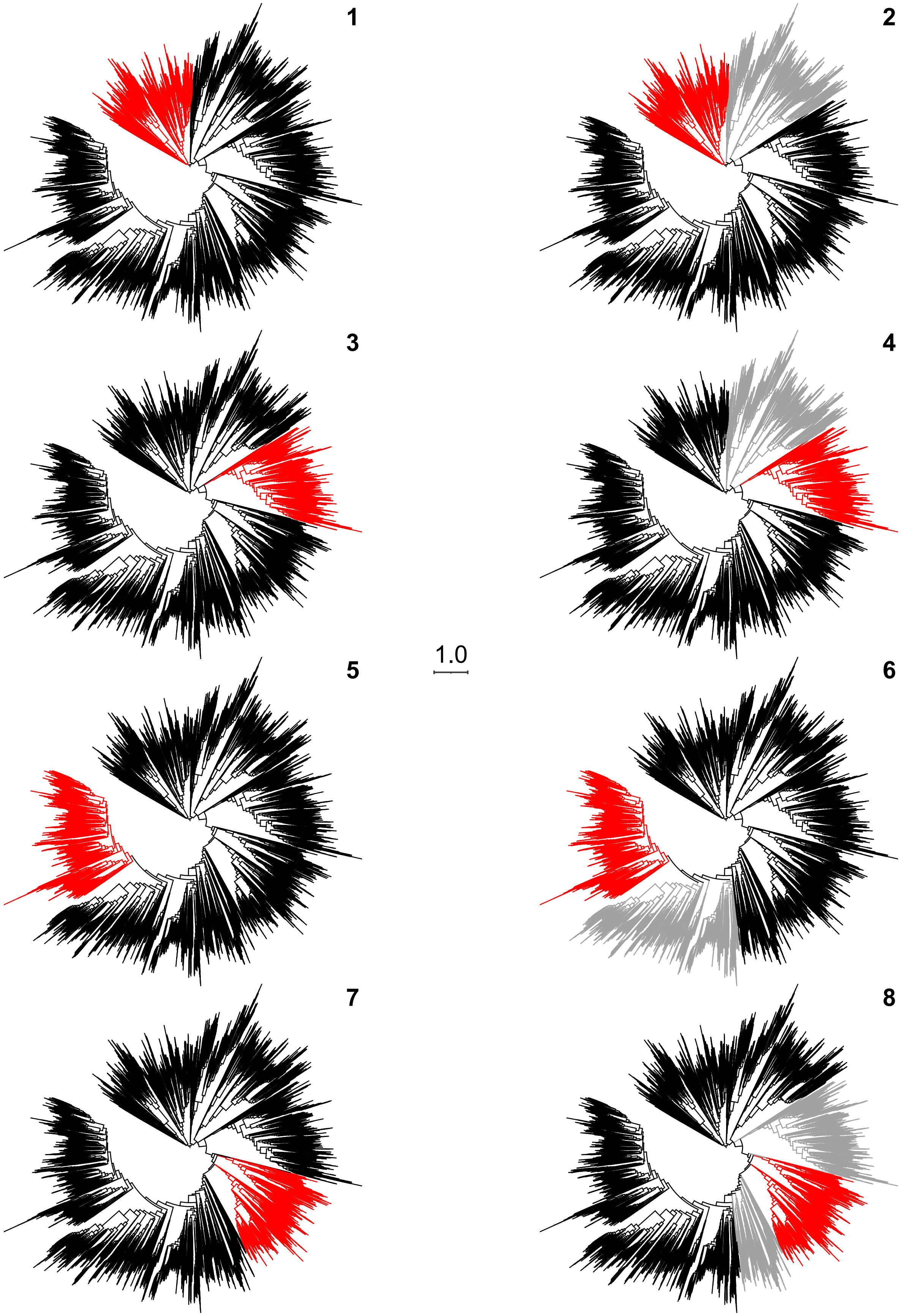

1
2
3
4
5
6
7
8
1.0

**Figure S8**. **Phylogenetically informed design of the data splits 1-8 of the portal protein dataset used for the training and testing of the models.** The phylogenetic tree relates all used portal proteins and is the same on all panels. For each data split, black branches show portal proteins used for the model training, red branches correspond to portal proteins used for the model testing, and gray branches refer to portal proteins that were not used neither in the model testing nor training. The phylogenetic tree is scaled with number of substitutions per site (see scale bar) and was rooted using the testing set of the split 1.

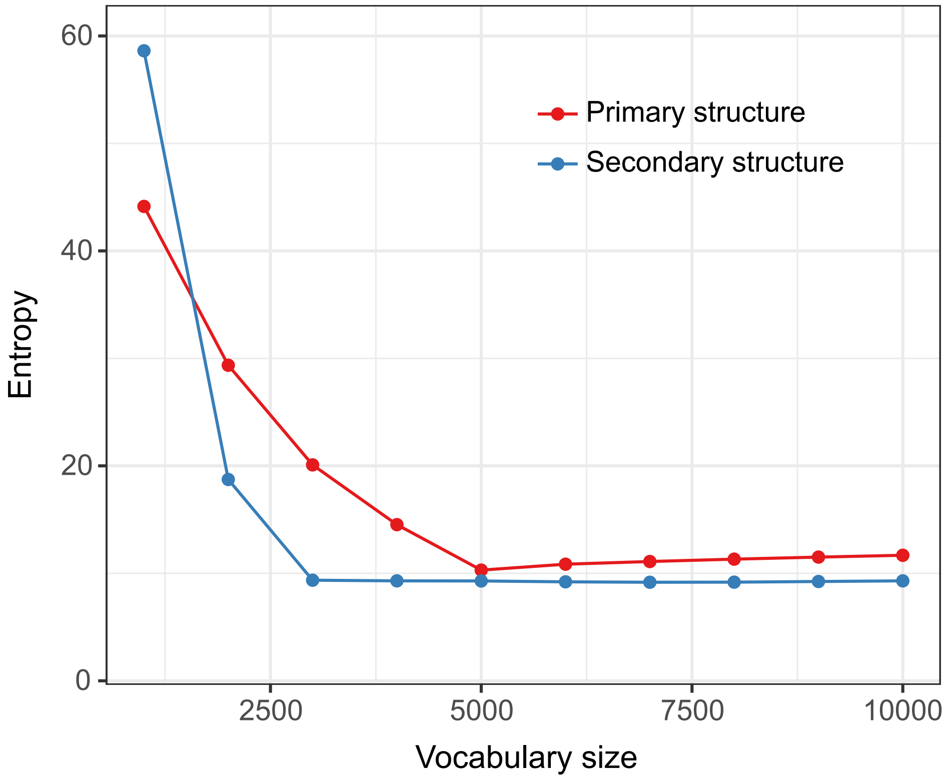


**Figure S9**. **Entropy values of models pre-trained on different vocabulary sizes.** The points correspond to the tested vocabulary sizes, which ranged from 1,000 to 10,000 tokens.
